# Activation of the CA2-vCA1 pathway reverses social discrimination dysfunction in *Shank3B* knockout mice

**DOI:** 10.1101/2022.03.28.486130

**Authors:** Elise C. Cope, Samantha H. Wang, Renée C. Waters, Betsy Vasquez, Elizabeth Gould

**Affiliations:** Princeton Neuroscience Institute, Princeton University, Princeton NJ 08544

## Abstract

Social memory dysfunction is a feature of several neuropsychiatric and neurodevelopmental disorders. Mutation or deletion of the *SHANK3* gene, which codes for a synaptic scaffolding protein, has been linked to autism spectrum disorder (ASD) and Phelan-McDermid syndrome, conditions associated with impairments in social memory. *Shank3B* knockout (KO) mice exhibit several behavioral abnormalities that may be analogous to symptoms of ASD, including social discrimination deficits. The CA2 region of the hippocampus integrates numerous afferents and sends a major output to the ventral CA1 (vCA1), circuitry that plays an important role in social memory. Despite finding few differences in excitatory afferents to the CA2 in *Shank3B* KO mice, chemogenetic activation of CA2 excitatory neurons restored social recognition function to wildtype (WT) levels. Specific activation of the CA2-vCA1 circuitry had a similar effect. Neuronal oscillations in the theta, gamma and, sharp wave ripple ranges have been linked to social memory, but we observed no differences in these measures between WT and *Shank3B* KO mice in the vCA1 in response to social stimuli. However, activation of CA2 excitatory neurons enhanced vCA1 theta power in *Shank3B* KO mice, concurrent with behavioral improvements. These findings suggest that stimulating adult circuitry in a mouse model with neurodevelopmental impairments can be sufficient to invoke latent function, particularly with respect to social memory dysfunction. The extent to which vCA1 network oscillations in the theta range are responsible for rescued behavioral function remains unknown.

## Introduction

Social memory is an important ability that gives rise to adaptive social interactions. Social memory dysfunction is a characteristic of several neuropsychiatric disorders, including autism spectrum disorder (ASD), schizophrenia, and major depressive disorder (Williams et al., 2005; Porcelli et al., 2019; Schafer and Schiller, 2019). Deficits in the ability to recognize individuals and to make associations between individuals and specific traits, emotions, events, and settings can significantly impair the formation and maintenance of social relationships (Kurtz et al., 2012; Garcia-Casal et al., 2017). These findings raise the importance of identifying the mechanisms of social memory as potential targets for therapeutic intervention.

Evidence from human studies indicates hippocampal involvement in social memory (Quian Quiroga, 2021; Montagrin et al., 2018), including reports of hippocampal abnormalities in individuals with conditions associated with social memory dysfunction (Laakso et al., 2000; Sheline et al., 2002; Brambilla et al., 2003; Kalmady et al., 2017). A growing number of studies aim to investigate mechanisms of social memory using experimental animals, with many reporting that the hippocampus plays an important role in this function (Kogan et al., 2000; Stevenson and Caldwell, 2014; Okuyama, 2018). Circuitry supporting social memory has been identified in the rodent hippocampus, with studies describing the CA2 as a social memory “hub” that integrates signals from a variety of afferents (Hitti and Siegelbaum, 2014). These afferents are both extrahippocampal, from the supramammillary nucleus and paraventricular nucleus of the hypothalamus, the cholinergic basal forebrain, and the entorhinal cortex, and also intrahippocampal, from the dentate gyrus and CA3 region. Taken together, the CA2 and these afferents play important roles in social recognition (Dudek et al., 2016; Raam et al., 2017; Piskorowski and Chevaleyre, 2018; Chiang et al., 2018; Chen et al., 2020; Lopez-Rojas et al., 2022).

Several types of network oscillatory patterns in the hippocampus have been associated with social behavior. Recent studies have shown that social stimuli result in changes to sharp wave ripples (SWRs), high frequency oscillatory events known to be associated with nonsocial memory consolidation and retrieval (Buzsáki, 2015; Joo and Frank, 2018), in both the CA2 (Oliva et al., 2020) and ventral CA1 (vCA1) (Rao et al., 2019). The CA2 communicates with the vCA1 (Meira et al., 2019; Brown et al., 2020), which serves as the main hippocampal output carrying social memory information (Okuyama et al., 2016; Watarai et al., 2021). Social stimuli also increase CA1 oscillations in the gamma and theta ranges (Villafranca-Faus et al., 2021; Brown et al., 2020), and several animal models of social dysfunction exhibit abnormal hippocampal gamma and theta power (Aoki et al., 2017; Arbab et al., 2018; Modi et al., 2019). Taken together, these findings suggest that atypical SWRs, as well as gamma and theta rhythms, in the CA2-vCA1 pathway may contribute to social memory impairment.

ASD represents a range of conditions with defining core symptoms (social impairments and restrictive interests/repetitive behaviors) of differing severity, as well as a wide range of potential comorbidities (intellectual disability, anxiety disorders, epilepsy) (NIMH, 2018). Perhaps not surprisingly, given the wide range of symptom presentation, the etiology of ASD appears to be multifactorial (Chaste and LeBoyer, 2012; Lai et al., 2014). It is generally accepted that ASD arises through a complex interaction between genes and the environment (Chaste and Leboyer, 2012). Genome wide association studies have identified over a hundred genes linked to ASD, with a significant number playing a role in neuronal communication (Satterstrom et al., 2020).

Due to this multifactorial etiology with clustered risk genes, understanding the mechanisms of social memory and associated dysfunctional neuronal circuitry may reveal translatable discoveries beyond genetic approaches. Among the genes shown to be linked to ASD is *SHANK3*, whose mutation or deletion also causes Phelan-McDermid syndrome, a condition that presents with similar symptoms to ASD (Costales and Kolevzon, 2015; DeRubeis et al., 2018). The *SHANK3* gene encodes the SH3 and multiple ankyrin repeat domains 3 protein, a synaptic scaffolding protein that functions to anchor receptors and ion channels to the postsynaptic site (Monteiro and Feng., 2017). Shank3 protein also plays a role in excitatory synapse and dendritic spine formation (Roussignol et al., 2005). Several *Shank3* models have been created in nonhuman primates and rodents using transgenic and CRISPR technologies. These models, which involve mutations or knockouts of parts or all of the *Shank3* gene, exhibit behavioral phenotypes analogous to the core symptoms of ASD, including social deficits and excessive repetitive behaviors (Zhou et al., 2019; Guo et al., 2019; Mei et al., 2016; Dhamne et al., 2017; Qin et al., 2018). These findings suggest that studies using *Shank3* animal models have the potential to provide translationally relevant information about circuits impacted in ASD and, importantly, to suggest points of intervention for improving function.

In the majority of cases, ASD symptoms persist throughout life (Blumberg et al., 2016). Studies have shown that adults with ASD experience lower life satisfaction (Griffiths et al., 2019; Moss et al., 2017; van Heijst and Geurts., 2015; Ayres et al., 2018) and are more likely to be unemployed and socially isolated than peers without ASD, due to the persistence of social impairments and socially disruptive behaviors (Orsmond et al., 2013; Moss et al., 2017; Roux et al., 2017; Griffiths et al., 2019; Solomon, 2020). These findings emphasize the need to identify interventions for adults with ASD. Along these lines, recent studies have shown that deep brain stimulation of the striatum diminishes excessive repetitive behavior and improves social communication in adults with ASD (Doshi et al., 2019; Davis et al., 2021). These promising results raise the possibility that activating brain regions involved in social memory might restore this ability as well. To investigate this in a mouse model of social dysfunction, we first confirmed that *Shank3B* knockouts (KO) have deficient social discrimination abilities, in that they respond similarly to novel and familiar mice, but that most excitatory afferents to the CA2 are similar to their wildtype (WT) littermates. We next used chemogenetics to activate excitatory neurons in the CA2 region, and also more directly the CA2-vCA1 pathway, which restored social discrimination abilities in adult *Shank3B* KO mice. We found that *Shank3B* KO mice have typical rhythms in the SWRs, gamma, and theta ranges in the vCA1, yet DREADD-induced improvement of social discrimination was accompanied by a boost in theta power.

## Results

### *Shank3B* KO mice have impaired social discrimination, but intact object location memory

We first confirmed previous reports that *Shank3B* KO mice have impaired sociability as well as impaired social discrimination memory (Peca et al., 2011; Tao et al., 2022). We utilized a three-trial direct social interaction test, with each trial separated by 24 hr, in which mice were exposed to a novel mouse (Novel 1) in trial 1, re-exposed to the same mouse (Familiar) in trial 2, and exposed to a second novel mouse (Novel 2) in trial 3 (Figure 1A). Adult WT mice typically prefer novelty and thus investigate novel mice more than familiar mice. To assess whether there were sex difference on genotype, we carried out a three-way ANOVA (sex x genotype x trial) and found no significant interaction between sex and genotype (*p* = 0.7246), so male and female data were collapsed for all subsequent analyses. We found the WT mice to have statistically significantly higher interaction times for the novel 1 and novel 2 than for the familiar mouse (Figure 1B, Supplementary Table 1). *Shank3B* KO mice did not have significant differences in interaction times across trials (Figure 1B, Supplementary Table 1) and had lower magnitude difference scores than WT mice (Figure 1C, Supplementary Table 1), indicating lower preference or discrimination of novelty. They also had lower novel interaction times as compared to WT, perhaps indicating reduced sociability (Figure 1B, Supplementary Table 1).

**Figure 1.**
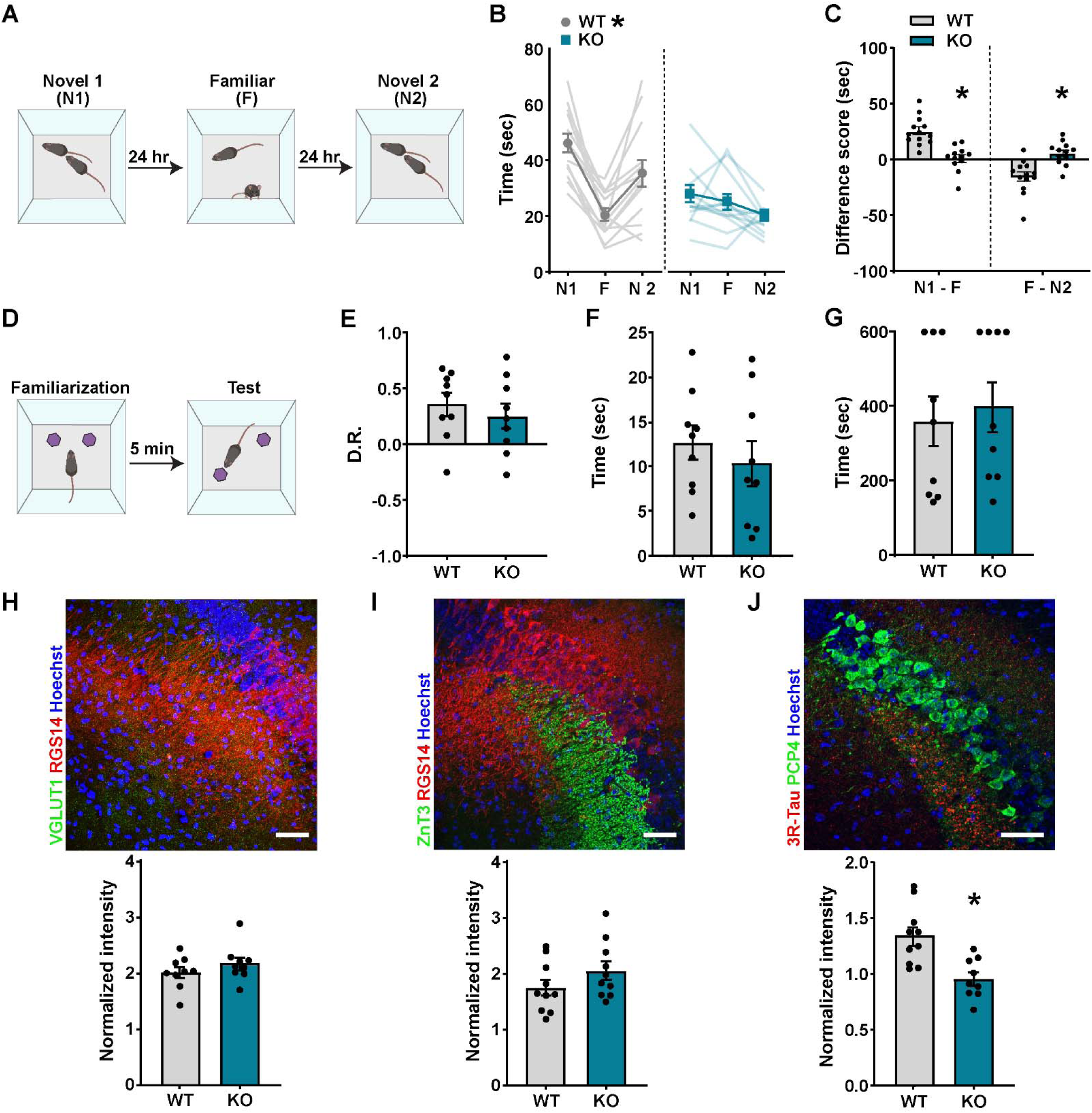
Compared to WT mice, *Shank3B* KO mice have impaired social, but not object location, discrimination, and mostly typical CA2 afferents. A) Schematic of social discrimination test. **B)** WT mice have lower interaction times for familiar mice (F) than first novel mice (N1) (p < 0.0001) and higher interaction times for second novel mice (N2) than F (*p* = 0.0112). *Shank3B* KO mice do not show different interaction times for F compared to N1 (*p* > 0.9999) or N2 (p = 0.1911). **C)** Compared to WT mice, *Shank3B* KO mice showed significant differences in difference scores (N1 minus F: *p* < 0.0001; F minus N2: *p* < 0.000491). For both comparisons (N1 minus F, F minus N2), *Shank3B* KO mice had difference scores that were closer to zero than WT. (for panels B and C, n = 13 for WT and n = 12 for KO) **D)** Schematic of object location test. **E)** WT and *Shank3B* KO showed no difference in discrimination ratios (DR) in the object location test (*p* = 0.517). **F)** There was no difference between genotypes in total time interacting with the objects during the test phase (*p* = 0.476). **G)** There was no difference in the amount of time it took to reach the criterion for familiarization with the objects *(p* = 0.6783) (E - G, n=9 for each genotype). **H)** Top: Confocal image of the CA2 immunolabeled with CA2 marker RGS14 (red), excitatory afferent marker VGLUT1 (green), and counterstained with Hoechst (blue). Bottom: WT and *Shank3B* KO mice have similar intensity of VGLUT1+ afferents in the CA2 (*p* = 0.2809) (n = 9 for each genotype). **I)** Top: Confocal image from the CA2 immunolabeled with CA2 marker RGS14 (red), mossy fiber marker ZnT3 (green), and counterstained with Hoechst (blue). Bottom: WT and *Shank3B* KO mice have similar intensity of ZnT3+ afferents in the CA2 (*p* = 0.1789) (n = 10 for each genotype). **J)** Top: Confocal image from the CA2 immunolabeled with CA2 marker PCP4 (green), abGC afferent marker 3R-Tau (red), and counterstained with Hoechst (blue). Bottom: Compared with WT, *Shank3B* KO mice have lower intensity of 3R-Tau+ afferents in the CA2 (*p* = 0.002) (n = 10 for WT and n = 9 for KO). For panels H-J, scale bars = 50 µm. See Supplementary Table 1 for complete statistics. Error bars represent SEM. \**p* < 0.05.

We then examined whether this deficit generalized to other hippocampal-dependent cognitive tests, particularly without social memory components. Using the object location memory test (Figure 1D), which requires the hippocampus (Barker and Warburton, 2011), we found that *Shank3B* KO mice were similarly capable as WT mice at distinguishing between objects in a novel versus familiar location (Figure 1E, Supplementary Table 1). There was also no difference between genotypes in their total investigation times of the objects (Figure 1F, Supplementary Table 1) or time to reach the criterion for familiarization (Figure 1G, Supplementary Table 1). Taken together, these data show that while *Shank3B* KO mice have significant impairments in social memory, they show typical object location memory.

### *Shank3B* KO mice exhibit similar avoidance behavior to WT mice

To consider the possibility that reduced sociability in *Shank3B* KO mice is the result of high levels of general avoidance behavior, we examined behavior on the elevated plus maze (EPM) (Supplementary Figure 1A, Supplementary Table 2). Indicators of avoidance behavior are percentage of time and entries onto open arms. *Shank3B* KO mice showed no differences on these measures (Supplementary Figure 1B, C, Supplementary Table 2), which suggests that *Shank3B* KO mice do not display more avoidance behavior on the EPM relative to WT mice, which is consistent with some (Dhamne et al., 2017) but not all (Liu et al., 2021) previous findings. *Shank3B* KO mice did show significant differences with lower percentage of time spent in the center (Supplementary Figure 1B, Supplementary Table 2) and fewer entries into the closed arms, as compared to WT mice (Supplementary Figure 1C, Supplementary Table 2). These differences might be better explained by activity or decision-making differences rather than avoidance behavior.

### Most afferents to CA2 appear similar in WT and *Shank3B* KO mice

Within the hippocampus, Shank3 protein is present in excitatory synapses that co-label with the vesicular glutamate transporter 1 (VGLUT1) (Heise et al., 2016). VGLUT1 labels axons from CA3 and entorhinal cortex pyramidal cells (Balschun et al., 2010; Zander et al., 2010; Heise et al., 2016), as well as from dentate gyrus granule cells (Heise et al., 2016). Since the CA3, entorhinal cortex, and dentate gyrus each directly project to the CA2 (Kohara et al., 2014; Middleton and McHugh, 2020) and each have been linked to social memory (Chiang et al., 2018; Leung et al., 2018; Lopez-Rojas et al., 2022), we analyzed intensity of VGLUT1 labeling in the CA2 and found no difference between WT and *Shank3B* KO mice (Figure 1H, Supplementary Table 1). Because Shank3 protein is especially concentrated in mossy fibers (Heise et al., 2016), we examined intensity of zinc transporter 3 (ZnT3), which labels mossy fibers and their synaptic vesicles, in the CA2 and also found no difference between WT and *Shank3B* KO mice (Figure 1I, Supplementary Table 1).

Next, because adult-born granule cells (abGCs) also project to the CA2 (Llorens-Martin et al., 2015) and have been linked to social memory (Cope et al., 2020), we analyzed the intensity of 3R-Tau, a microtubule-associated protein that labels abGC cell bodies and their mossy fibers (Llorens-Martin et al., 2012), in the CA2. Compared to WT mice, we found that *Shank3B* KO mice have lower 3R-Tau+ mossy fiber intensity in the CA2 (Figure 1J, Supplementary Table 1).

Since fewer afferents from abGCs to the CA2 might result from lower numbers of abGCs in the dentate gyrus of *Shank3B* KO mice, we then examined the number of 3R-Tau labeled cell bodies in the dorsal dentate gyrus and found a slight but significantly lower density in the suprapyramidal, but not infrapyramidal, blade of *Shank3B* KO mice (Supplementary Figure 2, Supplementary Table 2).

We next considered projections from extrahippocampal structures to the CA2 region, including glutamatergic inputs from the hypothalamic supramammillary nucleus (SUM) and cholinergic inputs from the medial septum, both of which have been implicated in social discrimination (Chen et al., 2020; Pimpinella et al., 2021; Robert et al., 2021). We examined the intensity of vesicular glutamate transporter 2 (VGLUT2), which is known to label SUM afferents to the CA2 (Heise et al., 2016), and vesicular acetylcholine transporter (VAChT), which is known to label cholinergic afferents (Al-Onaizi et al., 2017), and found that intensity of both markers was similar in the CA2 between WT and *Shank3B* KO mice (Supplementary Figure 3, Supplementary Table 2).

Our results suggest that social discrimination dysfunction in *Shank3B* KO mice is not the result of obvious abnormalities in several developmentally-generated inputs to this brain region, although an adult-generated afferent population appears to be diminished. The extent to which diminished abGC numbers and axons arise directly from *Shank3B* KO and contribute to social memory dysfunction remains unknown.

### Increasing CA2 activity improves social discrimination in *Shank3B* KO mice

Using chemogenetics, we tested whether activating excitatory neurons in CA2 would improve social discrimination in *Shank3B* KO mice. Excitatory DREADD virus (pAAV-CaMKIIa- hM3D(Gq)-mCherry) or control virus (pAAV-CaMKIIa-GFP) was bilaterally injected into the CA2 of WT and *Shank3B* KO mice (Figure 2A,B). Two weeks after surgery, mice were tested for social discrimination abilities using a direct social investigation test. Each virus-injected mouse was tested with both vehicle and CNO in separate behavior testing. For each behavioral testing bout, vehicle or CNO was injected 30 min prior to each novel and familiar mouse exposures.

**Figure 2.**
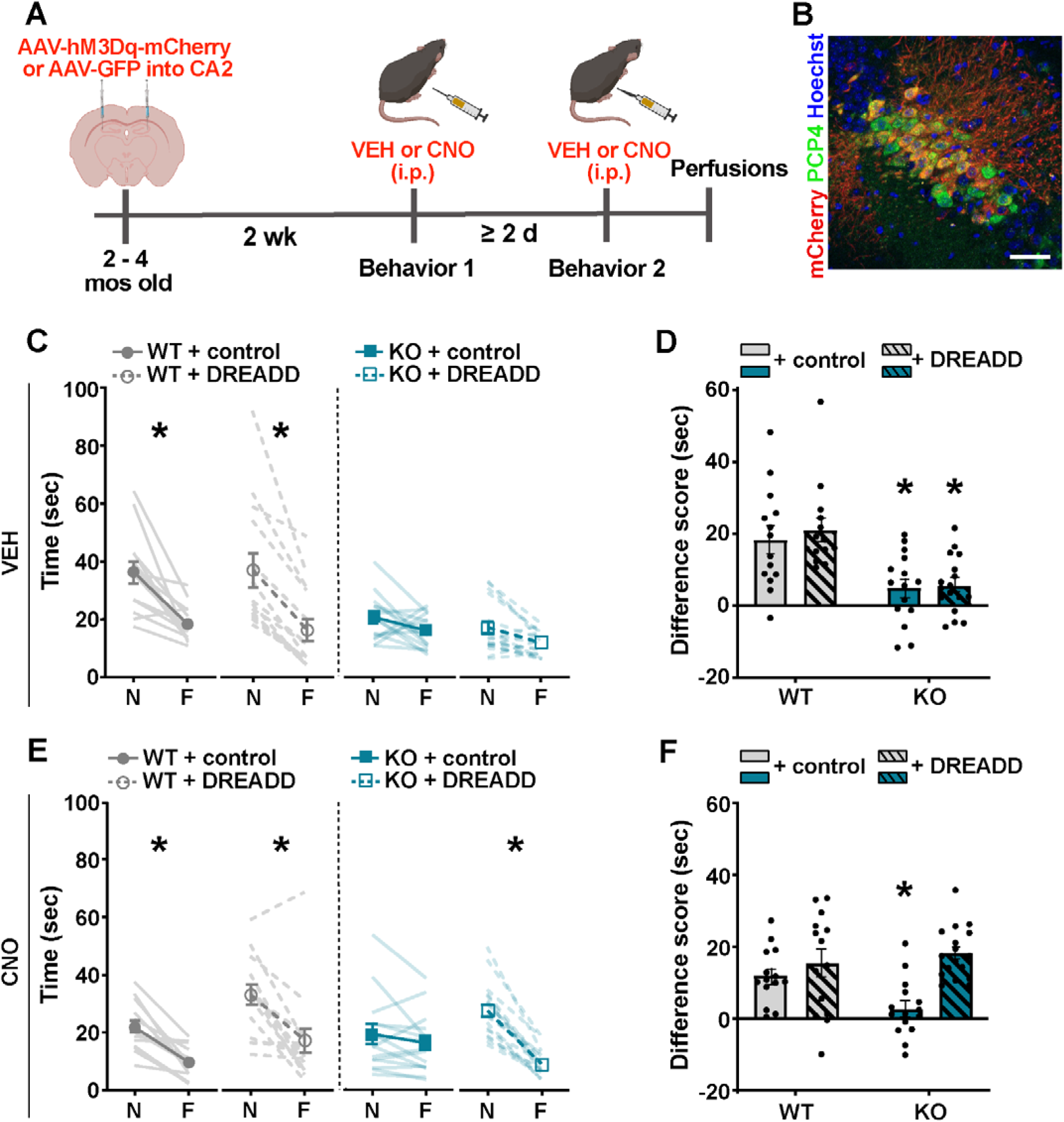
Chemogenetic activation of excitatory neurons in CA2 improves social discrimination in *Shank3B* KO mice. **A)** Timeline for virus infection, i.p. drug administration, and behavior testing. **B)** Confocal image from the CA2 immunolabeled with CA2 marker PCP4 (green), Gq AAV mCherry (red), and counterstained with Hoechst (blue) showing localized virus infection in CA2 neurons. Scale bar = 50 µm. **C)** Following vehicle (VEH) injections, virus- infected WT mice had greater interaction times for novel mice (N) than familiar mice (F) (WT + control virus, *p* < 0.0001; WT + DREADD virus, *p* < 0.0001), while virus-infected *Shank3B* KO mice showed no difference in interaction times of N compared to F (KO + control virus, *p* > 0.9999; KO + DREADD virus, *p* = 0.5679). **D)** Following VEH injections, WT virus groups had higher difference scores (N minus F) than *Shank3B* KO virus groups (WT control virus *vs* KO control virus: *p* = 0.0106; WT DREADD *vs* KO DREADD: *p* = 0.0014) **E)** Following CNO injections, virus-infected WT mice had greater interaction times for N than F (WT + control virus, *p* = 0.0002; WT + DREADD virus, *p* < 0.0001). Control virus-infected *Shank3B* KO mice had no difference in interaction times between N and F (*p* > 0.9999), while DREADD virus-infected *Shank3B* KO mice had greater interaction times for N than F (*p* < 0.0001). **F)** Following CNO injections, WT virus groups had higher difference scores than *Shank3B* KO control virus- infected mice (WT control virus *vs* KO control virus: *p* = 0.0777; WT DREADD virus *vs* KO control: *p* = 0.0039) but not *Shank3B* KO DREADD virus-infected mice (WT control virus *vs* KO DREADD virus: *p* = 0.4375; WT DREADD virus *vs* KO DREADD virus: *p* > 0.9999) and *Shank3B* KO control virus-infected mice had lower difference scores than *Shank3B* KO DREADD virus-injected mice (KO control virus *vs* KO DREADD virus: *p* = 0.0002) (C-F, n = 14 for WT + control virus or DREADD virus, n = 15 for KO + control virus, and n = 17 for KO + DREADD virus). See Supplementary Table 1 for complete statistics. Error bars represent SEM. \**p* < 0.05

Testing with vehicle and CNO were counterbalanced to control for any order effects with a minimum of two days between each test to allow for wash out of CNO (Figure 2A).

Consistent with our previous results from unoperated mice (Figure 1), in the vehicle trials, WT mice showed typical social discrimination, whereas, *Shank3B* KO mice showed impaired social discrimination. WT mice virus groups (control and DREADD virus) injected with vehicle showed lower interaction times for familiar mice than novel mice; whereas, KO mice virus groups (control and DREADD virus) injected with vehicle showed no such change across trials (Figure 2C, Supplementary Table 1). In addition, the difference in time investigating novel and familiar mice was relatively unchanged for *Shank3B* KO mice virus groups and was significantly different than WT mice virus groups (Figure 2D, Supplementary Table 1).

As expected, WT mice injected with either control or DREADD virus decreased their interaction times for familiar mice when administered CNO with no difference noted between control and DREADD virus groups (Figure 2E, Supplementary Table 1). CNO had no effects on social discrimination in *Shank3B* KO mice injected with control virus, while *Shank3B* KO mice injected with DREADD virus showed significantly greater investigation times with novel mice compared to familiar mice. *Shank3B* KO DREADD virus mice treated with CNO had social discrimination abilities that did not differ from WT mice (Figure 2E, Supplementary Table 1). Moreover, for the CNO injections, the difference in time investigating novel and familiar mice was not significantly different between the WT control and DREADD virus groups and the *Shank3B* KO DREADD virus group, while the *Shank3B* KO mice control virus group was significantly lower (Figure 2F, Supplementary Table 1).

### Increasing activity in the CA2 to vCA1 pathway improves social discrimination in *Shank3B* KO mice

Silencing the connection between the CA2 and ventral CA1 (vCA1) has been shown to produce social memory deficits in controls (Meira et al., 2018), including reducing the difference in interaction times with novel and familiar mice. We next explored whether activating CA2 afferents to vCA1 would improve social discrimination in *Shank3B* KO mice. WT and *Shank3B* KO mice received bilateral injections of excitatory DREADD virus (pAAV-CaMKIIa-hM3D(Gq)- mCherry) or control virus (pAAV-CaMKIIa-GFP) in the CA2 and were then implanted with bilateral cannulas into vCA1 to allow for local and selective excitation of CA2 projecting neurons (Figure 3A,B). Two weeks after surgery, mice were tested for social discrimination abilities using a direct social investigation test. Each virus-injected mouse was tested with both vehicle and CNO in separate behavior testing. For each behavioral testing bout, vehicle or CNO was infused into the vCA1 bilaterally 30 min prior to each novel and familiar mouse exposures. Testing with vehicle and CNO was counterbalanced to control for any order effects with a minimum of two days between each test to allow for wash out of CNO (Figure 3A).

**Figure 3.**
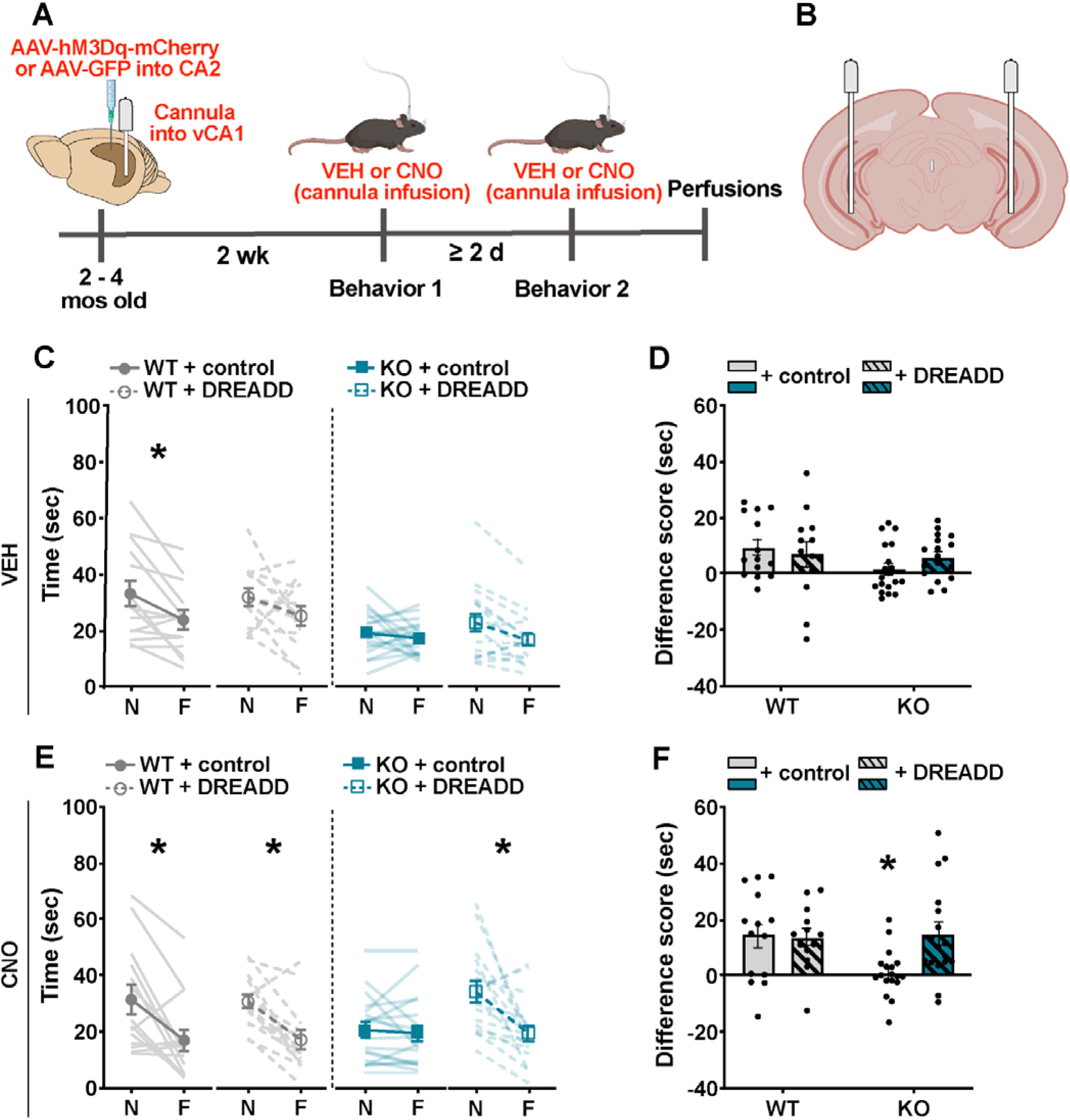
Chemogenetic activation of CA2 excitatory neurons projecting to the vCA1 improves social discrimination in *Shank3B* KO mice. A) Timeline for virus infection, cannula drug administration, and behavior testing. **B)** Schematic of the cannula placement sites in vCA1. **C)** Following vehicle (VEH) infusions into the vCA1, virus-infected *Shank3B* KO mice showed no significant difference in interaction times between novel mice (N) and familiar mice (F) (KO + control virus, *p* > 0.9999; KO + DREADD virus, *p* = 0.3094), while control virus- or DREADD virus-infected WT mice showed a decrease in interaction times between N and F, although the difference in WT+DREADD virus was not statistically significant (WT + control virus, *p* = 0.0270; WT + DREADD virus, *p* = 0.3566). It is worth noting that there was no effect of virus group (see Supplementary Table 1). **D)** Following VEH infusions into the vCA1, both WT virus groups had greater difference scores (N minus F) than *Shank3B* KO virus groups, although these comparisons were not statistically significant (*p* > 0.05, see Supplementary Table 1). **E)** Following CNO infusions into vCA1, WT virus groups and DREADD virus-injected *Shank3B* KO mice had lower interaction times of F than N (KO + DREADD, *p* = 0.0004; WT + control, *p* = 0.0021; WT + DREADD, *p* = 0.0071), while control virus-injected *Shank3B* KO mice did not (*p* > 0.9999). **F)** Following CNO infusions into vCA1, DREADD virus-injected *Shank3B* KO mice had higher difference scores (N minus F) than control virus-injected *Shank3B* KO mice (p=0.0268) and no difference from WT mice injected with control virus (p>0.9999) or DREADD virus (p>0.9999) (n = 14 for WT + control, n = 13 for WT + DREADD, n = 18 for KO + control, n = 17 for KO + DREADD). See Supplementary Table 1 for complete statistics. Error bars represent SEM. \**p* < 0.05

Following vehicle infusions into vCA1, compared to their interaction times with novel mice, WT control virus mice showed typical social discrimination as evidenced by their lower interaction for familiar mice, whereas *Shank3B* KO control virus mice showed no difference in their interaction times for familiar mice (Figure 3C, Supplementary Table 1). Compared to WT mice, the difference in time investigating novel and familiar was on average lower for *Shank3B* KO mice, but the difference between WT and *Shank3B* KO did not reach statistical significance (Figure 3D, Supplementary Table 1).

CNO infusions into the vCA1 had no effect on WT control virus or WT DREADD virus mice as demonstrated by their typical social memory (Figure 3D, Supplementary Table 1). As expected, *Shank3B* KO control virus mice infused with CNO into the vCA1 had impaired social discrimination abilities (Figure 3D, Supplementary Table 1). However, *Shank3B* KO DREADD virus mice injected with CNO into the vCA1 displayed typical social discrimination (Figure 3D). After vCA1 CNO infusion, *Shank3B* KO control virus mice showed no change in time spent investigating novel and familiar mice with difference times that were significantly lower than WT control and *Shank3B* KO DREADD virus mice (Figure 3E, Supplementary Table 1). WT mice virus groups (control and DREADD virus) had lower interaction times for familiar mice than novel mice when infused with CNO, showing no difference between control and DREADD virus groups. *Shank3B* KO control virus group retained their lack of difference across trials even when infused with CNO, while the *Shank3B* KO mice DREADD virus group infused with CNO showed significantly greater investigation times with novel mice compared to familiar mice. The *Shank3B* KO DREADD virus group treated with CNO had social discrimination abilities that did not differ from WT virus groups treated with CNO (control and DREADD virus) (Figure 3E, Supplementary Table 1). Moreover, for the CNO infusions, the difference in time investigating novel and familiar mice was not significantly different between WT virus groups and the *Shank3B* KO DREADD virus group, while the *Shank3B* KO control virus group was significantly lower (Figure 3F, Supplementary Table 1).

### vCA1 theta power does not differ between WT and *Shank3B* KO mice, but increases in *Shank3B* KO mice after chemogenetic activation of the CA2

vCA1 theta (4-12 Hz) power is associated with higher avoidance behavior (Adhikari et al., 2010; Padilla-Coreano et al., 2019; Murthy et al., 2019) and because compared to WT mice, *Shank3B* KO mice spend considerably less time investigating novel mice, we explored whether *Shank3B* KO mice had higher vCA1 theta power during exposure to novel mice. WT and *Shank3B* KO mice received bilateral injections of excitatory DREADD virus (pAAV-CaMKIIa-hM3D(Gq)- mCherry) or control virus (pAAV-CaMKIIa-GFP) in the CA2 and were then implanted with recording electrodes into vCA1 (Figure 4A). We found no difference in vCA1 theta power between WT and *Shank3B* KO mice (Figure 4C, Supplementary Table 1), nor did we find a correlation between vCA1 theta power and the time spent investigating novel mice in WT or *Shank3B* KO mice (Pearson’s rank correlation coefficient test, WT: *r* = -0.2965, *p* = 0.4384; KO: *r* = 0.1291, *p* = 0.7405). Taken together with our EPM results, these findings suggest that low sociability observed in *Shank3B* KO mice is not directly related to general avoidance, or to avoidance-associated vCA1 theta power.

**Figure 4.**
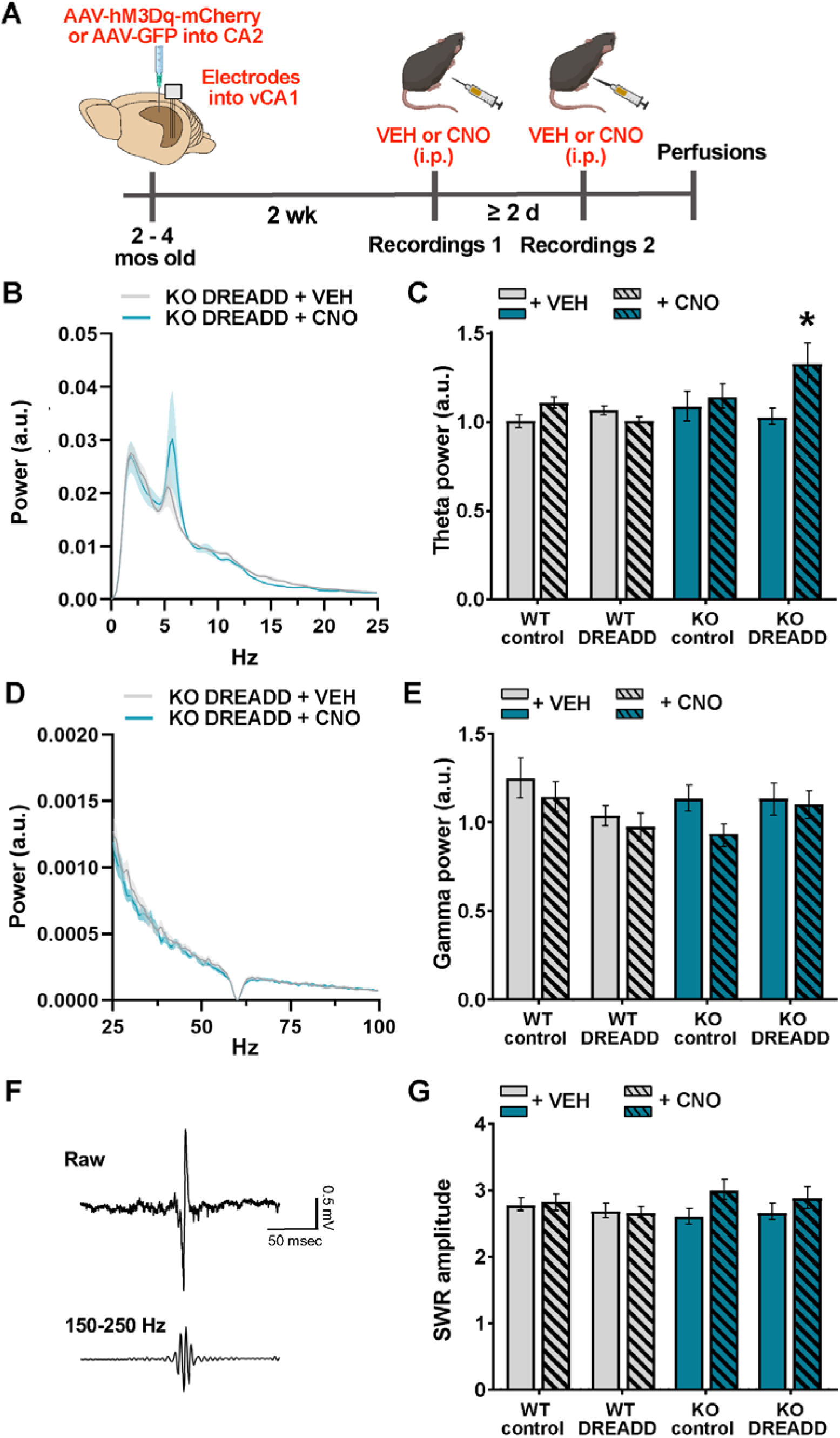
Chemogenetic activation of the CA2 increases vCA1 theta power in *Shank3B* KO mice, but does not affect gamma power nor SWRs. A) Timeline for virus infection, electrode placement, and neural recordings. **B)** vCA1 power spectra during exposure to a novel mouse (0-25 Hz) of vehicle (VEH)- and CNO-treated *Shank3B* KO DREADD virus-injected mice. Shaded area represents SEM. **C)** vCA1 theta power (4-12 Hz) was not different between WT and *Shank3B* KO virus groups treated with VEH (*p* > 0.05, see Supplementary Table 1). *Shank3B* KO DREADD mice treated with CNO had higher theta power than *Shank3B* KO DREADD virus-injected mice treated with VEH (*p* = 0.0147), while no other groups had significant differences. **D)** vCA1 power spectra during exposure to a novel mouse (25-100 Hz) of VEH- and CNO-treated *Shank3B* KO DREADD virus-injected mice. Shaded area represents SEM. **E)** vCA1 low gamma power (30-55 Hz) was not different between WT and *Shank3B* KO virus groups injected with VEH or CNO during novel mouse exposure (*p* > 0.05, see Supplementary Table 1). **F**) Representative example of SWRs (Top: Raw LFP trace. Bottom: Filtered SWR trace) recorded in the vCA1. **G)** There were no differences in normalized SWR amplitude after exposure to a familiar mouse between WT and *Shank3B* KO groups control virus- or DREADD virus-infected groups treated with VEH, nor was this changed by CNO- induced activation of the CA2 (*p* > 0.05, see Supplementary Table 1) (C, E, and G, n = number of electrodes, 9-10 mice per group with 3 electrodes for each mouse). See Supplementary Table 1 for complete statistics. Error bars represent SEM. \**p* < 0.05

Because social stimuli have been shown to increase hippocampal theta power (Villafranca- Faus et al., 2021), and because chemogenetic activation of CA2 and the CA2-vCA1 pathway increased investigation of novel, but not familiar, mice (Figure 2E, 3E), we examined whether *Shank3B* KO CA2 DREADD-infected mice treated with CNO showed changes in vCA1 theta power during novel mouse exposure. We found that chemogenetic activation of the CA2 increased vCA1 theta power in the *Shank3B* KO DREADD virus group, but not in the WT groups (Figure 4C, Supplementary Table 1).

### vCA1 gamma power does not differ between WT and *Shank3B* KO mice, and remains unchanged after chemogenetic activation of the CA2

Previous studies have shown that low gamma (30-55 Hz) is altered by chemogenetic activation of CA2 neurons in the CA2/proximal dCA1 (Alexander et al., 2016). Thus, we investigated whether DREADD activation of CA2 neurons altered low gamma power in WT and *Shank3B* KO mice during exposure to novel mice and found no differences across any groups (Figure 4D,E, Supplementary Table 1). The difference between our findings and the previous report may be due to recording from different areas of the CA1 (dorsal in Alexander at al., 2016 versus ventral in the present study).

### SWRs are similar between WT and *Shank3B* KO mice, and not changed with chemogenetic activation of CA2

SWRs have been linked to consolidation and retrieval of spatial memories (Joo and Frank, 2018). The vCA1 has been shown to have increased SWR events in the presence of a social stimulus (Rao et al., 2019) and CA2-generated SWRs, which are known to influence vCA1 SWRs, are necessary for social memory (Oliva et al, 2020). Given that our behavioral findings suggest diminished function in the CA2-vCA1 pathway in *Shank3B* KO mice, we explored the possibility of alterations in SWRs in the vCA1 that may be rescued by chemogenetic activation of the CA2. Since SWRs typically only occur during sleep and behavioral immobility (Joo and Frank, 2018), we first measured SWR frequency, with SWR number as a function of immobility time. However, *Shank3B* KO mice spend significantly more time immobile than WT mice (WT: 100.8 +/- 10.18 sec, KO: 184.8 +/- 10.29 sec, unpaired t-test, t_17_ = 5.785, *p* < 0.0001), and reduced SWR frequency in *Shank3B* KO mice (WT: 0.6367 +/- 0.05620 event frequency, KO: 0.3460 +/- 0.01708 event frequency, unpaired t-test, t_49_ = 4.703, *p* < 0.0001), an effect that clearly confounds the SWR frequency analysis. To eliminate this confound, we focused our analysis on SWR number within the first minute of exposure to a familiar mouse, a time frame when retrieval of social memories likely occurs. We detected SWR events in the vCA1 during exposure to familiar mice (Figure 4G, Supplementary Table 1). We found no difference in SWR amplitude, duration, or number between WT or *Shank3B* KO mice (Figure 4G; Supplementary Figure 4A,B, Supplementary Table 1,2) or following chemogenetic activation of the CA2 (Figure 4G; Supplementary Figure 4A,B, Supplementary Table 1,2). These findings suggest that although WT and *Shank3B* KO mice differ in their ability to discriminate between novel and familiar mice, there is no difference in SWR measures in response to a familiar mouse, nor is there any notable change in response to CA2 activation.

## Discussion

Our findings suggest that *Shank3B* KO mice have impaired social discrimination, with a major deficit in time spent investigating novel social stimuli, compared to WT littermates. We found no difference between groups in a simple object location task or in avoidance behavior, suggesting that *Shank3B* KO mice have some unimpaired non-social memory and that differences in sociability seem unlikely due to general avoidance. Because Shank3 is a synaptic scaffolding protein, we next investigated whether multiple developmentally-generated afferents to the CA2 are different in *Shank3B* KO mice compared to WT mice. Unexpectedly, we found that markers of CA2 afferents known to be involved in social discrimination, including VGLUT1, VGLUT2, ZnT3, and VAChT, were not different from WT in *Shank3B* KO mice. In contrast, we found that 3R-Tau, a marker of abGC axons, was diminished in the CA2 of *Shank3B* KO compared to WT mice. Because the CA2 has been associated with social memory, we next attempted to improve social discrimination by chemogenetically activating excitatory neurons in the CA2 of *Shank3B* KO mice. We found that activating the CA2 with DREADD virus + CNO restored social discrimination in *Shank3B* KO mice to WT levels. We next investigated whether specific activation of projections from the CA2 to the vCA1 would also produce this effect and found restored social discrimination in *Shank3B* KO mice to WT levels. These findings suggest that *Shank3B* KO social discrimination ability can be improved by activating the CA2-vCA1 pathway in adulthood. We next tested whether vCA1 SWRs, which have been linked to memory consolidation and retrieval, differ between WT and *Shank3B* KO mice and found similarities in amplitude, number, and duration, as well as no change in vCA1 SWRs during chemogenetic manipulation. Although we also detected no differences in vCA1 theta or gamma power in *Shank3B* KO compared to WT mice for control virus or vehicle treatments, theta power was increased in *Shank3B* KO mice infected with DREADD virus and treated with CNO beyond WT levels. Increased vCA1 theta power was observed during exposure to the novel mouse, which is the experience that elicited behavioral change after CA2 activation in *Shank3B* KO mice.

Our behavioral data suggest that low social investigation of novel mice by *Shank3B* KO mice is likely not the result of greater social avoidance compared to WT mice. However, we observed no evidence of high avoidance behavior of *Shank3B* KO mice on a non-social task, the elevated plus maze, findings that are consistent with some (Dhamne et al., 2017), but not all (Liu et al., 2021), published reports. We also found no evidence of increased vCA1 theta power, an electrophysiological correlate of non-social avoidance behavior (Padilla-Coreano et al., 2019), in *Shank3B* KO mice displaying low levels of novel social investigation (*Shank3B* KO control virus, *Shank3B* KO DREADD virus + vehicle), and no correlation between investigation times of novel mice and vCA1 theta power. In fact, the one experimental group that showed elevated vCA1 theta power (*Shank3B* KO DREADD virus + CNO) had been subjected to a manipulation that increased investigation of novel mice compared to familiar mice by *Shank3B* KO mice. Taken together, these results suggest that low investigation times are not the result of global behavioral inhibition. Low investigation of a novel social stimulus may represent a number of specific deficits, including faulty recognition of a novel stimulus as familiar, inattention to social stimuli, reduced motivation/reward associated with social stimuli, and/or inability to recognize social novelty, all of which have been reported in humans with ASD (Guillory et al., 2021; Hedger and Chakrabarti, 2021; Aldridge-Waddon et al., 2020; Weigelt et al., 2012; Chevallier et al., 2012). More specifically related to the *Shank3B* mouse model, individuals diagnosed with both Phelan-McDermid syndrome and ASD exhibit reduced social attention and reduced social novelty recognition under certain conditions (Guillory et al., 2021).

Social novelty recognition, along with broader social memory, has been linked to the CA2 region in mice (Middleton and McHugh, 2020). Electrophysiological studies have shown increased firing of CA2 pyramidal cells in response to a novel social stimulus (Donegan et al., 2020). Additional studies have identified afferents to the CA2 as being involved in these functions. In particular, projections from the SUM and the cholinergic medial septum have been linked to novelty recognition (Chen et al., 2020; Wu et al., 2021; Pimpinella et al., 2021), whereas those from the vasopressinergic paraventricular nucleus, the entorhinal cortex, and the dentate gyrus seem more likely associated with memory of familiar social stimuli (Smith et al., 2016; Lopez-Rojas et al., 2022; Cope et al., 2020). Despite the fact that Shank3, a synaptic scaffolding protein, is expressed in the hippocampus, we found no differences between WT and *Shank3B* KO mice in markers of several of these afferents to the CA2. These results were unexpected and suggest that compensation for the lack of Shank3 might occur during development, particularly at synapses with high concentrations of this molecule, including those made by mossy fibers, as well as those expressing VGLUT1 (Heise et al., 2016). In this regard, it may be relevant that Shank3 colocalizes with other synaptic scaffolding proteins (Shank1 and 2) (Heise et al., 2016), which may enable the development of functional synapses in the absence of Shank3. Despite the lack of global abnormalities in CA2 afferents of *Shank3B* KO mice, we did observe a difference in the afferents from abGCs in the dentate gyrus. abGCs, like mature granule cells, are known to project to the CA2 (Kohara et al., 2014; Llorens-Martin et al., 2015) and have been shown to participate in social memory (Cope et al., 2020). Although it is possible that the lower numbers of abGCs and their afferents to CA2 contribute to social discrimination deficits in *Shank3B* KO mice, it seems unlikely to be the main contributor, since transgenic reduction of abGCs reduces social memory function without affecting investigation times of novel social stimuli (Cope et al., 2020).

Regardless of the lack of global abnormality of several excitatory afferent labels in CA2 of *Shank3B* KO mice, chemogenetic activation of CA2 pyramidal cells in general or the CA2-vCA1 pathway directly in these mice restored social discrimination to resemble that of WT mice. Analysis of social investigation times suggests that the main effect of CA2 excitatory neurons or CA2-vCA1 activation is to increase investigation times of the novel mouse. This effect was only seen in the *Shank3B* KO mice, with no changes observed in the WT mice following identical treatment. These findings suggest that an, as yet, unidentified dysfunction upstream or within the vCA1, potentially of the CA2 and/or CA2-vCA1 pathway, exists in *Shank3B* KO mice, but that stimulating existing circuitry is sufficient to invoke latent function.

Oscillatory activity in the CA2 and vCA1 regions has been linked to social memories. CA2 SWRs have been causally associated with social memory consolidation (Oliva et al., 2020), raising the possibility that abnormalities in their number, magnitude or duration might contribute to social memory dysfunction in *Shank3B* KO mice. Because previous studies have shown that SWRs are also involved in retrieval of nonsocial memories (Joo and Frank, 2018), we hypothesized that vCA1 SWRs might differ between WT and *Shank3B* KO mice during familiar mouse presentation, a time when social memory retrieval is most likely. However, we found no differences in multiple SWR measures between WT and *Shank3B* KO mice, as well as no changes in these measures after chemogenetic activation of CA2 excitatory neurons. It remains possible that vCA1 SWRs might differ between WT and *Shank3B* KO mice but that the parameters of our recordings did not capture this effect. Along these lines, it is relevant to note that a recent study found diminished SWR amplitude in *Shank3B* KO compared to WT mice during the 2 hr rest period after social interaction, a time likely important for social memory consolidation (Tao et al., 2022). However, since the main difference we noted in social interaction between WT and *Shank3B* KO mice was with investigation of novel social stimuli, which does not involve memory consolidation or retrieval, it seems likely that this social novelty deficit is not dependent on differences in SWRs.

Hippocampal theta and gamma rhythms have been associated with novelty recognition (Penley et al., 2013; Kitanishi et al., 2015; Zheng et al., 2016; Brown et al., 2020; Kragel et al., 2020; Park et al., 2021; Wang et al., 2021). Since social stimuli have been shown to increase hippocampal theta and gamma oscillations (Tendler and Wagner, 2015; Villafranca-Faus et al., 2021; Brown et al., 2020) and mouse models of social dysfunction have been associated with alterations in hippocampal oscillations at both frequencies (Arbab et al., 2018; Aoki et al., 2017; Cheaha and Kumarnsit, 2015), we examined theta and gamma power in the vCA1 of WT and *Shank3B* KO mice in the presence of a novel mouse. Unexpectedly, we found no differences in either theta or gamma power in the absence of CA2 activation. Chemogenetic activation of CA2, however, produced an increase in vCA1 theta, but not gamma, power in *Shank3B* KO mice. This effect was not observed in WT mice, paralleling the results of our behavioral studies. Taken together, these findings suggest that although theta power does not appear to be atypical in *Shank3B* KO mice in response to novel mice, it is increased under conditions associated with a restoration of novel mouse investigation to WT levels. Thus, although the underlying abnormality of *Shank3B* KO mice in social behavior circuitry remains unknown, elevated vCA1 theta power corresponds with enhanced investigation of novel mice by *Shank3B* KO mice. The extent to which elevated vCA1 theta power is causally associated with behavioral changes remains unknown.

The possibility that chemogenetic activation of CA2-induced vCA1 theta power in *Shank3B* KO mice is responsible for increased investigation of novel social stimuli may seem contradictory with reports of causal links between vCA1 theta power and avoidance behavior (Padilla-Correano et al., 2019), however, that association has only been observed with nonsocial avoidance, and we found no correlation between theta power and social investigation times in WT or *Shank3B* mice without DREADD virus and CNO. Studies have shown that the vCA1 is heterogeneous anatomically and functionally, with subsets of neurons connected to the hypothalamus/medial prefrontal cortex, amygdala, and nucleus accumbens, each serving different functions, including avoidance, learning, and social behavior respectively (Gergues et al., 2020). Indeed, theta coherence between hippocampus and different target regions has been reported to differ after exposure to social stimuli versus nonsocial stimuli, with the latter linked to defensive/avoidance behavior (Tendler and Wagner, 2015). Thus, it seems likely that chemogenetic activation of the CA2 may activate theta oscillations among subsets of vCA1 neurons that have downstream connections with regions associated specifically to social behavior. Along these lines, it is likely relevant that both vCA1 pyramidal neurons and parvalbumin-positive (PV+) interneurons, known targets of CA2 pyramidal neurons (Okuyama et al., 2016; Nasrallah et al., 2019), play important roles in social novelty detection and social discrimination (Deng et al., 2019). Since PV+ cells participate in the generation of theta rhythm (Amilhon et al., 2015), activating CA2 pyramidal cells may initiate theta oscillations among vCA1 PV+ interneurons, which produces rhythmic inhibition of specific populations of vCA1 pyramidal neurons. Studies have shown that inhibition of vCA1 pyramidal cells by PV+ interneurons is necessary for the recognition of social novelty (Nasrallah et al., 2019), and, conversely, that overactivation of vCA1 pyramidal cells reduces investigation of novel mice (Okuyama et al., 2016). Taken together with the present results, these findings suggest that *Shank3B* KO mice may have deficient vCA1 PV+ cell function, which can be overcome by stimulation of CA2 afferents and the initiation of greater vCA1 theta power. However, since we detected no differences in vCA1 theta power between WT and *Shank3B* KO mice without CA2 activation, any abnormality in vCA1 PV+ cell function likely involves functions other than the ability to generate theta oscillations.

Collectively, our findings suggest that *Shank3B* KO mice have deficits in social discrimination as a result of reduced investigation of novel social stimuli. Despite the lack of gross morphological abnormality in the CA2 of *Shank3B* KO mice, we found that chemogenetic stimulation of the CA2 and the CA2-vCA1 circuit was sufficient to restore social investigation of novel stimuli. Behavioral restoration in *Shank3B* KO mice with CA2 activation were associated with increased vCA1 theta power, but not gamma power or alterations in SWRs. These findings suggest that activation of a hippocampal social memory circuit in adulthood is sufficient to restore a behavioral deficit arising from a neurodevelopmental genetic anomaly.

## Methods

### Animals

All animal procedures were approved by the Princeton University Institutional Animal Care and Use Committee and were in accordance with the National Research Council Guide for the Care and Use of Laboratory Animals. To generate WT and *Shank3B* KO mice, adult male and female *Shank3B*+/- (JAX Stock no. 17688) mice were obtained from The Jackson Laboratory and bred at Princeton University using a heterozygous X heterozygous strategy. At postnatal day 15, mice were genotyped by Transnetyx using real-time PCR. Mixed groups of male and female WT and male and female KO mice were used as test animals for these experiments and HET mice that were the same sex and age as the test mice were used as stimulus animals. All mice were group housed by genotype and sex in Optimice cages on a reverse 12/12 h light/dark cycle. For social memory and object location memory behavioral studies, as well as histological studies, 6- to 8-week old mice were used (n=9-13/group). For elevated plus maze studies, 2-5 month old mice were used (n=20/group). For behavioral experiments involving chemogenetic activation of the CA2 and CA2-vCA1 pathway, 2-5 month old mice were used (n=13-18/group). For electrophysiological experiments, 2-5 month old mice were used (n=9-10/group).

### Surgical procedures

Mice were deeply anesthetized with isoflurane (2-3%) and placed in a stereotaxic apparatus (Kopf) under a temperature controlled thermal blanket. The head was levelled using bregma, lambda, and medial-lateral reference points before craniotomy was performed. Each mouse received bilateral injections (15 nl/ hemisphere at a rate of 15 nl/min) of either excitatory DREADD virus pAAV-CaMKIIa-hM3D(Gq)-mCherry (Addgene Cat# 50476-AAV5, titer: 1.7x10^13^) or control virus pAAV-CaMKIIa-EGFP (Addgene Cat# 50469-AAV5, titer: 4.3x10^12^) into the CA2 using the following coordinates from Bregma: -1.82 AP, +/-2.15 ML, and -1.67 DV. The virus was delivered using a 10 µl syringe with a 33-gauge beveled needle (NanoFil) controlled by a microinjection pump (WPI). The needle remained in place for an additional 5 min after the injection was completed to prevent backflow of the virus upon removal.

For activating the CA2-vCA1 projections, mice injected with DREADD or control virus into the CA2 were implanted with a cannula guide extending 4 mm (Plastics One, Cat# C315GS-5/SP) into the vCA1 (-3.5 AP, +/-3.45 ML). Dummy cannulas (Plastics One, Cat# C315DCS-5/SPC) were inserted into the guides and the guide was lowered to -3.8 mm. Cannula guides were kept in place using metabond and dental cement (Bosworth Trim).

For vCA1 recordings, mice injected with control or DREADD virus into the CA2 were implanted unilaterally with a custom-made 3 wire electrode array (Microprobes) into the right hemisphere of the vCA1 (electrode 1, AP: -3.3, ML: 3.45, DV: -3.8 with each electrode separated by 200 nm). Four bone screws were implanted on the skull and one screw was used as a ground. A ground wire was wrapped around the ground screw and covered with metallic paint. Electrode implants were kept in place using Metabond and dental cement (Bosworth Trim). Two to four weeks after surgeries, mice were i.p injected or cannula infused with either clozapine-N-oxide (CNO) or vehicle (see CNO administration) before being tested on behavioral tasks (see Behavioral assays) and/or underwent electrophysiological recordings (see Electrophysiology recordings).

### CNO administration

Each CA2 virus-injected mouse underwent social discrimination testing (with novel and familiar stimulus mice) twice, once after CNO i.p. injection or vCA1 cannula infusion and once after vehicle i.p. injection or vCA1 cannula infusion. The order of drug administration (CNO or vehicle) was counterbalanced across groups. Because previous studies have shown that DREADD manipulations of neurons are transient and return to baseline by 10-24 hours post-CNO injections (Alexander et al., 2009; 2018; Ray et al., 2011), mice were given a minimum 2- day rest period between CNO and vehicle tests. 30 min prior to both the novel (trial 1) and familiar (trial 2) stimulus mouse exposure, test mice received CNO or vehicle i.p. injections or vCA1 cannula infusions. For systemic administration of CNO, CA2 virus injected mice received i.p. injections of 1 mg/kg of CNO (dissolved in DMSO and then suspended in saline) or vehicle. To activate the CA2-vCA1 pathway, CA2 virus injected mice with implanted cannula received cannula infusions of CNO into the vCA1 under light isoflurane anesthesia (2%). After the dummy cannula was removed, an internal cannula projecting 0.8 mm (Plastics One, Cat# C315IS-5/SPC) from the tip of the guide cannula was inserted. 1 μl of CNO (2 μg/μl of CNO dissolved in DMSO and then suspended in saline) (Chang and Gean, 2019) or vehicle was infused per hemisphere over 1 minute into the vCA1 using a syringe pump (Harvard apparatus) mounted with a 1 µl syringe (Hamilton). The internal cannula remained in place for 1 additional minute after the infusion was completed to allow for diffusion of the drug. Mice were returned to their home cages and resumed typical ambulatory activity from the light anesthesia within 5 min.

### Behavioral assays

All behavioral testing occurred during the active cycle for mice (dark cycle). The testing arena was an open field box or an elevated plus maze (see below for details).

#### Social discrimination memory testing

To assess social discrimination, two versions of the direct social interaction test were adapted from previously established protocols (Hitti and Siegelbaum, 2014; Smith et al., 2016; Cope et al., 2020; 2021). Each version of this test consisted of two or three social stimulus trials, each separated by 24 hr. For behavioral characterization studies, each mouse was tested once with a three social stimulus trials paradigm. For virus manipulation studies, each mouse underwent social discrimination testing two times, each time with either vehicle or CNO treatment (order of injections or infusions was counterbalanced across groups) with vehicle and CNO treatment separated by at least a 2-day wash-out period. Prior to the test beginning, mice were acclimated to the behavior testing room for at least 30 min prior to the first social stimulus trial. The testing was conducted in low lighting in an open-field box (23 x 25 x 25 cm). For the three-social stimulus trial paradigm, the test mouse and a never-before-encountered mouse (Novel 1, trial 1) were placed together in the open field box and allowed to interact for 5 min. After this interaction period, the test mouse was returned to their home cage for 24 hr and then placed back into the open field arena with the previously encountered mouse (Familiar, trial 2). 24 hours after the second trial, the test mouse was introduced to a new, novel mouse (Novel 2, trial 3). For experiments involving virus manipulations, mice underwent two social stimulus trials during which the test mouse was introduced to a novel mouse (Novel 1) in trial 1, followed by the same novel mouse (Familiar) in trial 2. Sex-matched non-littermate HET mice were used as stimulus mice for social discrimination testing. For each trial, the interaction time of the test mouse with the stimulus mouse was measured from video recordings. Social interaction was defined as direct sniffing of the stimulus mouse’s anogenital region or body, following, or allogrooming that was initiated by the test mouse.

#### Object location memory

To assess a form of non-social memory, the object location test was used (Barker and Warburton, 2011). The testing was conducted in low lighting in an open-field box (23 x 25 x 25 cm). For 5 min, twice per day, for 3 days, mice were habituated to the testing arena as previously described (Cope et al., 2018a, 2018b, 2021). Mice were habituated to the objects on the third day of habituations. Two objects, each <8cm in height and width, were placed in the testing arena in a random orientation. Different, but visually identical, objects were used for habituation and testing. On the testing day, mice underwent a familiarization trial followed by a test trial. In the familiarization trail, two identical objects were positioned on the same side of the testing box (6 cm away from the walls and 10 cm between each other). Mice were free to explore the objects until they reached 30s of cumulative exploration time with both objects or up to a maximum of 10 min elapsed. The criterion for object exploration was directing their nose at 2 cm or less distance from the object. After the familiarization trial, mice were placed in their home cage for 5 min. Between trials, one object (moved object) was rotated 180° and moved to the opposite wall of the chamber so that it was diagonal to the first object, while the other object was not moved (familiar object). The moved object was counterbalanced throughout testing. For the test trial, mice were returned to the testing arena and were free to explore the objects for 2 min. The object exploration times were scored manually from video recordings. A discrimination ratio (DR) was calculated for each mouse as follows: (Time exploring moved object – time exploring familiar object) / total time exploring objects).

#### Elevated plus maze

Mice were placed on an elevated plus maze that consisted of an elevated (50 cm) plus-shaped track with two arms that were enclosed with high walls (30 cm) and two open arms that had no walls and illuminated to 200 lux. All arms were 50 cm in length. During testing, the mouse was placed in the center of the maze and allowed to explore for 5 min. The number of entries into the open and closed arms and time spent in the open arms, closed arms, and center was measured for each mouse from video recordings.

### Electrophysiology recordings

Local field potentials (LFPs) were recorded using a wireless head stage (TBSI, Harvard Biosciences). Mice were habituated to the weight of recording headstage using a custom headstage with equivalent weight in the home cage for 5 min the day before the first test and in the open field arena for 5 min on day of testing. The two social stimulus trial paradigm was used. For each trial, 30 min after drug administration, the test mouse was placed with the stimulus mouse and LFPs recorded continuously for 5 min. To get a baseline measurement, LFPs were also recorded for 3 min in the open field box prior to exposure to the social stimulus each day. The data were transmitted to a wireless receiver (Triangle Biosystems) and recorded using NeuroWare software (Triangle Biosystems).

### Electrophysiological analyses

All recordings were analyzed using Neuroexplorer software (Nex Technologies). For theta and gamma analyses, continuous LFP data were notched at 60 Hz and band-pass filtered from 0 to 100 Hz. To normalize theta and gamma oscillations, the sum of power spectra values from 0 to 100 Hz were set to equal 1. To obtain power estimates within theta (4-12 Hz) and low gamma (30-55 Hz) bands, the summed power across time for the entire session within each frequency was taken. For SWR analyses, continuous LFP data were notched at 60 Hz and band-pass filtered from 150 and 250 Hz. Signals were then Hilbert transformed and z-scored. SWR events were defined as exceeding three standard deviations of a 15 ms rolling-average amplitude threshold, excluding deflections within 15ms of a previous ripple, and were detected using a custom Python code. To determine SWR event frequency, the number of SWRs detected were normalized to immobility time (defined as quiet wakefulness). The mean amplitude and duration of the detected SWRs were calculated across the recording session. Theta and gamma power were analyzed across the 5 min of Trial 1, during exposure to the Novel conspecific, and normalized by dividing by the baseline trial. SWR numbers, amplitude, and duration were analyzed across the first 1 min of Trial 2, during exposure to the Familiar conspecific, and normalized by dividing by the baseline trial.

### Histology

Mice were deeply anesthetized with Euthasol (Virbac) and were transcardially perfused with cold 4% paraformaldehyde (PFA). Extracted brains were post-fixed for 48 h in 4% PFA at 4°C followed by an additional 48 h in 30% sucrose at 4°C for cryoprotection before being frozen in cryostat embedding medium at -80°C. Hippocampal coronal sections (40 µm) were collected using a cryostat (Leica). Sections were blocked for 1½ hr at room temperature in a PBS solution that contained 0.3% Triton X-100 and 3% normal donkey serum. Sections were then incubated overnight while shaking at 4°C in the blocking solution that contained combinations of the following primary antibodies: mouse anti- three microtubule-binding domain tau protein (3R-tau, 1:500, Millipore, Cat# 05-803), rabbit anti-purkinje cell protein 4 (PCP4, 1:500, Sigma-Aldrich, Cat# HPA005792), rat anti-mCherry (1:1000, Invitrogen, Cat# M11217), mouse anti-regulator of G protein signaling (RGS14, 1:500, UC Davis/NIH NeuroMab, Cat# 75–170), rabbit anti-zinc transporter 3 (ZnT3, 1:500, Alomone labs, Cat# AZT-013), rabbit anti-vesicular glutamate transporter 2 (VGLUT2, 1:500, Synaptic Systems, Cat# 135 403), rabbit anti-vesicular glutamate transporter 1 (VGLUT1, 1:250, Invitrogen, Cat# 48-2400), rabbit anti-vesicular acetylcholine transporter (VAChT, 1:500, Synaptic Systems, Cat# 139 103). For 3R-Tau immunohistochemistry, sections were subjected to an antigen retrieval protocol that involved incubation in sodium citrate and citric acid buffer for 30 min at 80°C prior to blocking solution incubation. Washed sections were then incubated for 1½ hr at room temperature in secondary antibody solutions that contained combinations of the following secondaries: donkey anti-rat Alexa Fluor 568 (1:500, Abcam), donkey anti-mouse Alexa Fluor 568 or 647 (1:500, Invitrogen), or donkey anti-rabbit Alexa Fluor 488 (1:500, Invitrogen). Washed sections were then counterstained with Hoechst 33342 for 10 min (1:5,000 in PBS, Molecular Probes), mounted onto slides, and coverslipped with Vectashield (Vector labs). Slides were coded until completion of the data analysis.

### Histological analysis

#### Optical intensity measurements

Z-stack images of the CA2 and corpus callosum were taken on a Leica SP8 confocal using LAS X software and a 40x oil objective. The CA2 was defined by PCP4 or RGS14 labeling. Collected z-stack images were analyzed for optical intensity in Fiji (NIH). A background subtraction using a rolling ball radius (50 pixels) was applied to the image stacks. A region of interest (ROI) was drawn and the mean gray value was collected throughout the image stack. In the CA2, the ROI was confined to the stratum lucidum for 3R-Tau and ZnT3, the stratum radiatum and lacunosum moleculare for VGLUT1, and the pyramidal layer and stratum oriens for VGLUT2 and VAChT. The mean gray value of the ROI was calculated for each z-slice and the maximum mean gray value for each z-stack was taken. That maximum of the CA2 ROI was divided by the maximum of the corpus callosum ROI for each section. Each brain’s normalized intensity was the average of 3 sections.

#### Cell density measurements

The number of 3R-Tau+ cells were counted in the dorsal dentate gyrus of the hippocampus on 4 neuroanatomically matched sections using an Olympus BX-60 microscope with a 100x oil objective. The counts for the suprapyramidal blade and infrapyramidal blade of the dentate gyrus were analyzed separately and area measurements were collected using Stereo Investigator software (MBF). The density of 3R-Tau was determined by dividing the total number of positively labeled cells by the volume of the subregion (ROI area multiplied by 40 µm section thickness).

### Statistical analysis

For histological analyses, data sets were analyzed using an unpaired two-tailed Student’s t-test or Mann Whitney U tests. For behavioral analyses involving two group comparisons, data sets were analyzed using either unpaired two-tailed Student’s t-tests or a repeated measures two-way ANOVA, as appropriate. For behavioral analyses involving virus manipulations, data sets were analyzed using either a two-way ANOVA or a repeated measures three-way ANOVA, as appropriate. Bonferroni post hoc comparisons were used to follow up any significant main effects or interactions of the ANOVAs. Pearson’s correlation coefficient test was used to analyze the association between theta power and social investigation times. All data sets are expressed as the mean ± SEM on the graphs and statistical significance was set at p<0.05 with 95% confidence. GraphPad Prism 9.2.0 (GraphPad Software) or Excel (Microsoft) were used for statistical analyses. All graphs were prepared using GraphPad Prism 9.2.0 (GraphPad Software). Statistical values (n sizes, p values, and statistical test) are reported in the figure legends or supplementary tables.

## Acknowledgments

This work was supported by the National Institutes of Health, NIMH 1R01 MH118631-01 to EG. The authors thank Biorender for assistance with the figure schematics, Monica Hanani and Kristen A. Pagliai for assistance with histological analyses, and Blake J. Laham for helpful comments on the project and for providing the Python code for the sharp wave ripple analyses.

## Supplementary figures

**Supplementary Figure 1.**
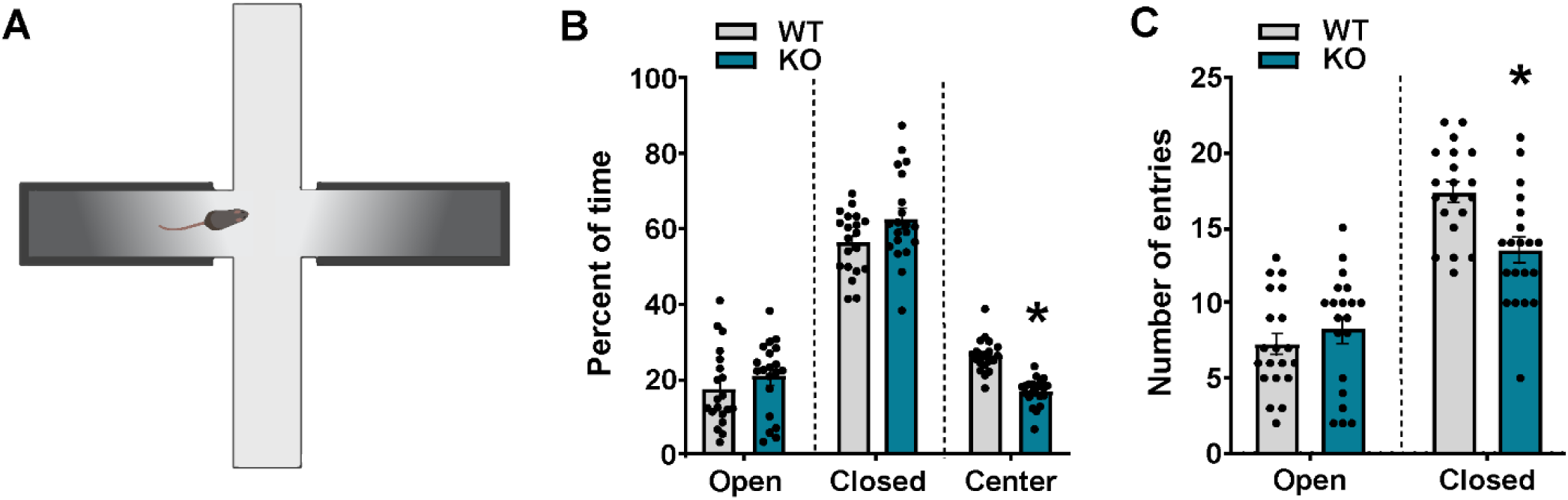
Compared to WT mice, *Shank3B* KO mice do not show increased avoidance behavior in the elevated plus maze. **A)** Schematic of the elevated plus maze test. **B)** Compared to WT mice, *Shank3B* KO mice showed no significant difference in percentage time in the open arms (*p* = 0.327) or closed arms (*p* = 0.0509), but a decrease in the percentage of time spent in the center (*p* < 0.0001). **C)** While the number of entries into the open arms was not different between genotypes (*p* = 0.406), *Shank3B* KO mice made fewer entries into the closed arms (*p* = 0.0011). \**p* < 0.05 compared to WT mice (n = 20 for each genotype), see Supplementary Table 2 for complete statistics. Error bars represent SEM.

**Supplementary Figure 2.**
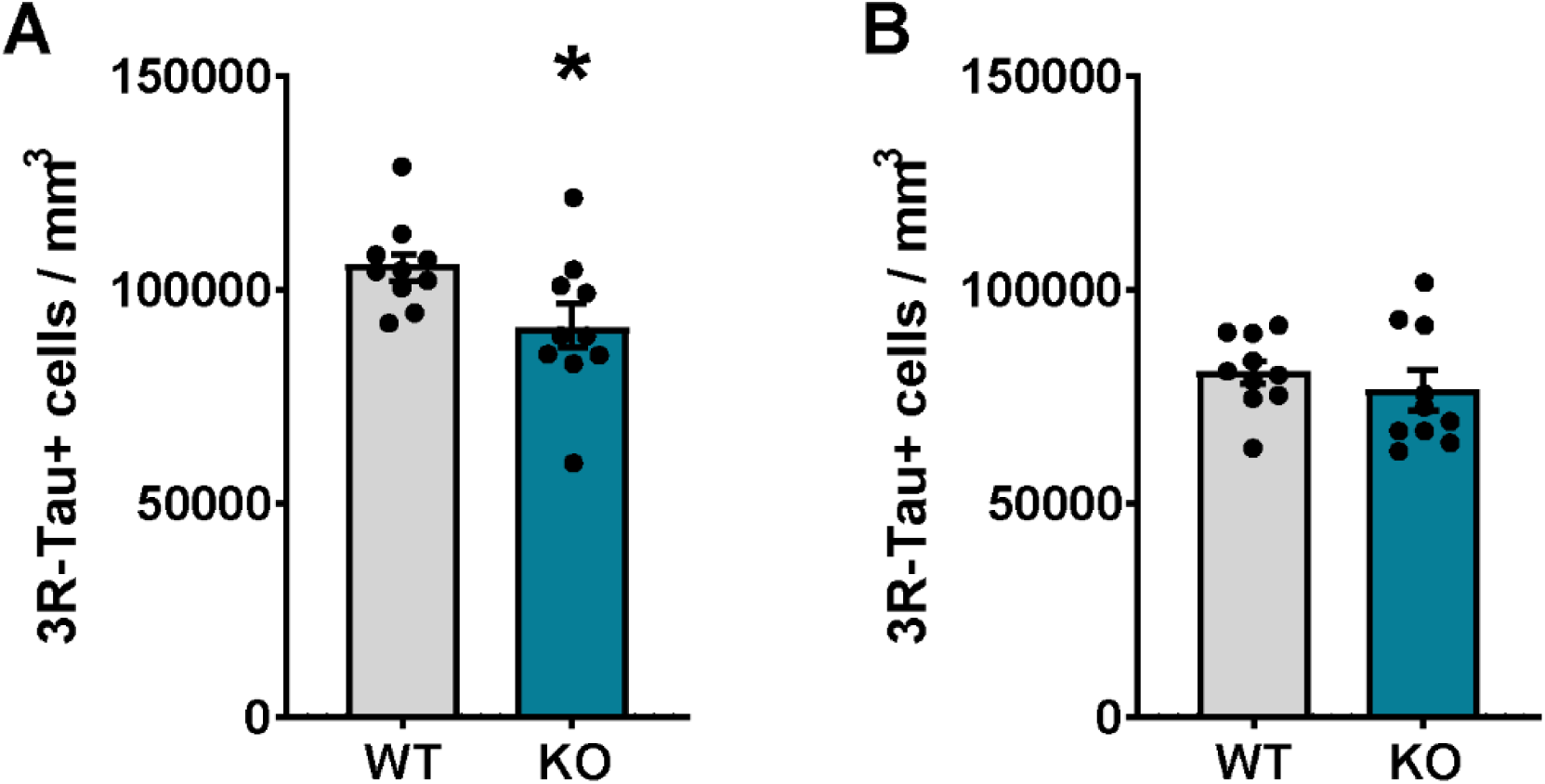
Compared to WT mice, *Shank3B* KO mice had a lower density of abGCs in the suprapyramidal, but not infrapyramidal, blade of the DG. **A)** Compared to WT mice, *Shank3B* KO mice had lower density of 3R-Tau+ cell bodies in the suprapyramidal blade (*p* = 0.0360), **B)** but not in the infrapyramidal blade (*p* = 0.416), of the DG. \**p* < 0.05 compared to WT mice (n = 10 per group), see Supplementary Table 2 for complete statistics. Error bars represent SEM.

**Supplementary Figure 3.**
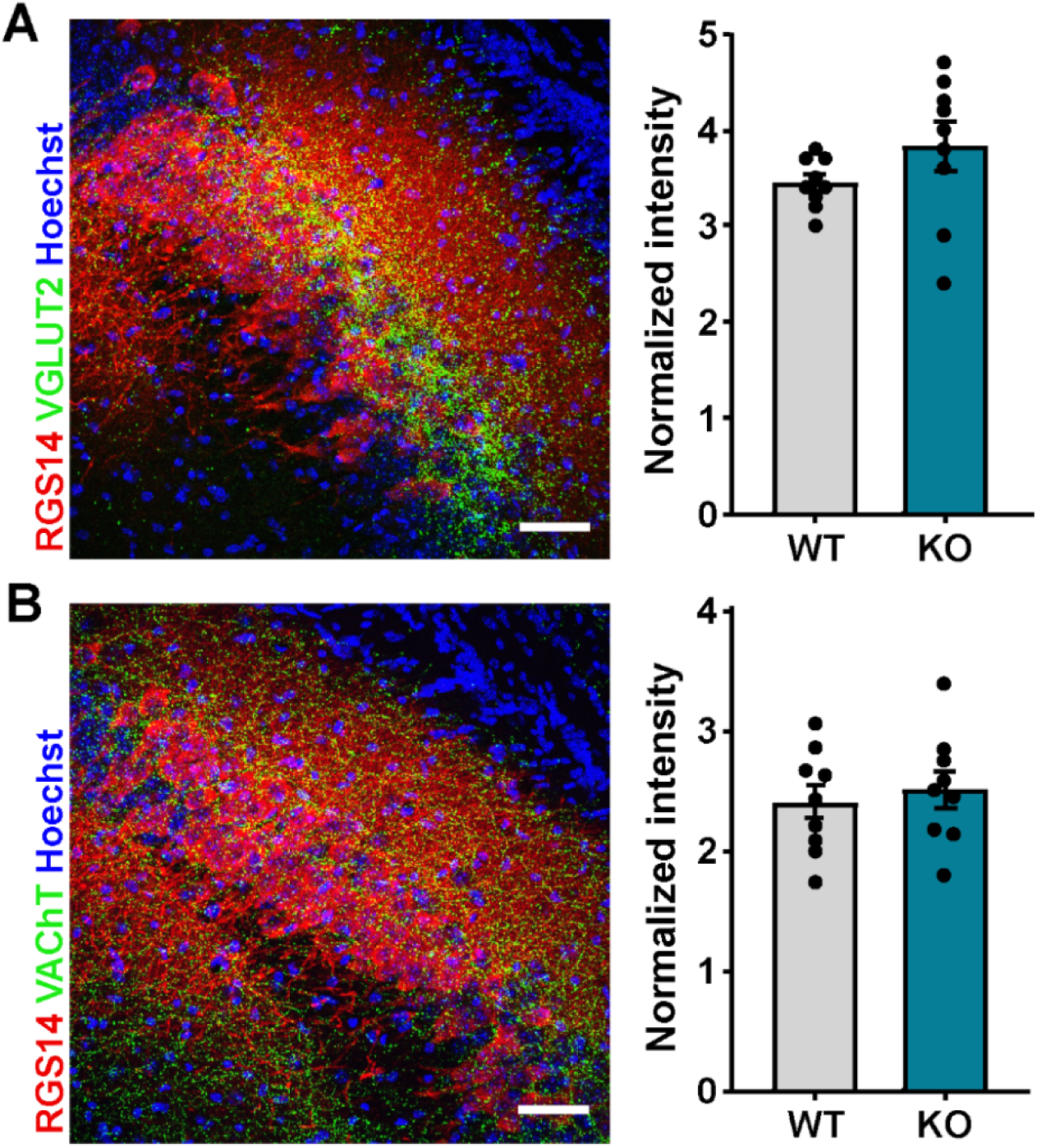
Compared to WT mice, *Shank3B* KO mice showed no difference in VGLUT2+ nor VAChT+ afferents in the CA2. **A)** Left: Confocal image from the CA2 immunolabeled with CA2 marker RGS14 (red), SUM afferent marker VGLUT2 (green), and counterstained with Hoechst (blue). Right: Compared to WT mice, *Shank3B* KO mice had similar intensity of VGLUT2+ afferents in the CA2 (*p* = 0.0984). **B)** Left: Confocal image from the CA2 immunolabeled with CA2 marker RGS14 (red), afferent marker VAChT (green), and counterstained with Hoechst (blue). Right: Compared to WT mice, *Shank3B* KO mice had similar intensity of VAChT+ afferents in the CA2 (*p* = 0.6202) (n = 9 for each genotype), see Supplementary Table 2 for complete statistics. Error bars represent SEM.

**Supplementary Figure 4.**
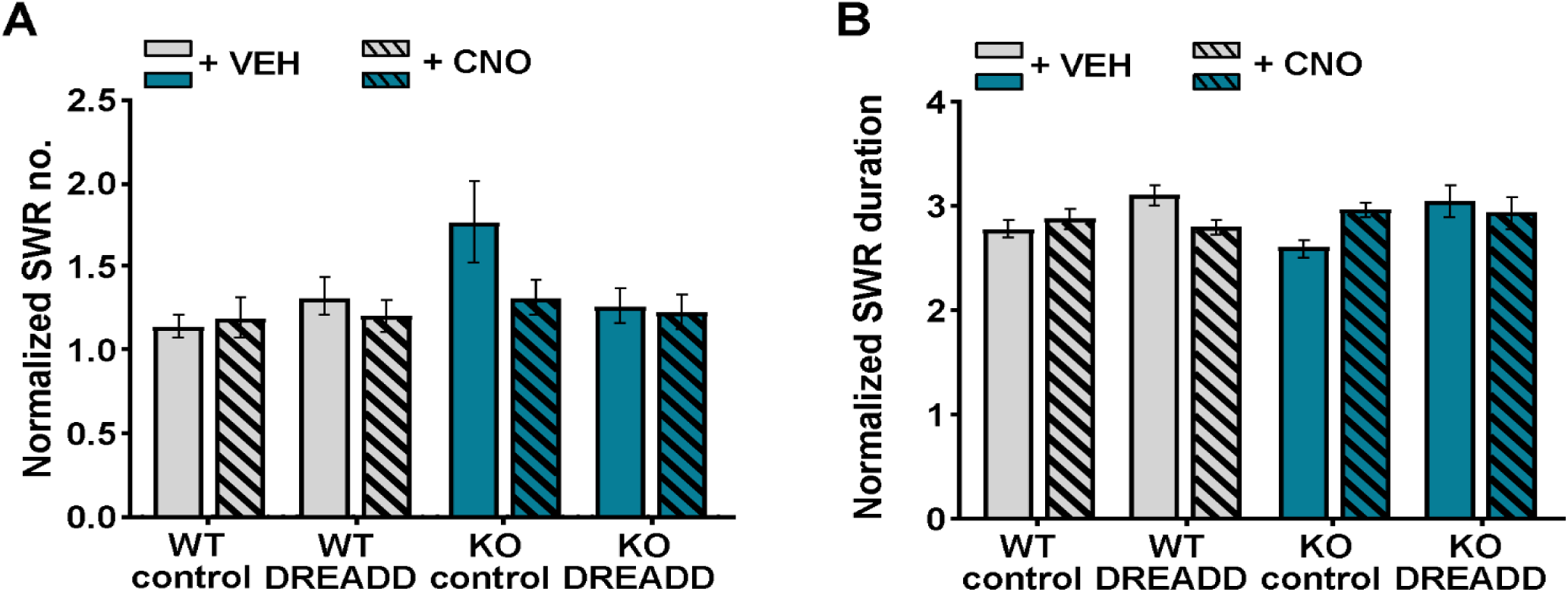
Compared to WT mice, *Shank3B* KO mice showed no difference in SWR numbers nor duration during familiar mouse exposure. **A)** During familiar mouse exposure, there was no difference in SWR numbers between genotypes, nor was this changed by CNO-induced activation of the CA2. **B)** During the first minute of familiar mouse exposure, there was no difference in SWR duration between genotypes, nor was this changed by CNO-induced activation of the CA2 (number of electrodes (9-10 mice per group with 3 electrodes for each mouse), see Supplementary Table 2 for complete statistics. Error bars represent SEM.

**Supplementary Table 1.** Detailed statistics for main figures.

| Data set | Type of test | Confidence |
| --- | --- | --- |
| Figure 1B | two-way repeated measures ANOVA with Bonferroni post hoc comparisons | <u>ANOVA</u><br>trial: $F_{(1.389,31.95)} = 10.33$, $p = 0.0012$<br>genotype: $F_{(1,23)} = 12.18$, $p = 0.002$<br>trial X genotype: $F_{(2,46)} = 9.358$, $p = 0.0004$<br><u>post hoc comparisons</u><br>WT N1 vs WT F: $p < 0.0001$<br>WT F vs WT N2: $p = 0.0112$<br>KO N1 vs KO F: $p > 0.9999$<br>KO F vs KO N2: $p > 0.1911$<br>WT N1 vs KO N1: $p = 0.0009$ |
| Figure 1C | unpaired t-test | N1 minus F: $t_{23} = 5.086$, $p < 0.0001$<br>F minus N2: $t_{23} = 4.055$, $p < 0.000491$ |
| Figure 1E | unpaired t-test | $t_{16} = 0.662$, $p = 0.517$ |
| Figure 1F | unpaired t-test | $t_{16} = 0.730$, $p = 0.476$ |
| Figure 1G | unpaired t-test | $t_{16} = 0.4225$, $p = 0.6783$ |
| Figure 1H | unpaired t-test | $t_{16} = 1.116$, $p = 0.2809$ |
| Figure 1I | unpaired t-test | $t_{18} = 1.399$, $p = 0.1789$ |
| Figure 1J | unpaired t-test | $t_{17} = 3.647$, $p = 0.002$ |
| Figure 2C | three-way repeated measures ANOVA with Bonferroni post hoc comparisons | <u>ANOVA</u><br>trial: $F_{(1,56)} = 75.00$, $p < 0.0001$<br>genotype: $F_{(1,56)} = 215.88$, $p = 0.0002$<br>virus: $F_{(1,56)} = 0.7528$, $p = 0.3893$<br>trial X genotype: $F_{(1,56)} = 25.95$, $p < 0.0001$<br>trial X virus: $F_{(1,56)} = 0.3769$, $p = 0.5417$<br>genotype X virus: $F_{(1,56)} = 0.4959$, $p = 0.4842$<br>trial X genotype X virus: $F_{(1,56)} = 0.1461$, $p = 0.7037$<br><u>post hoc comparisons</u><br>WT control virus N vs F: $p < 0.0001$<br>WT DREADD virus N vs F: $p < 0.0001$<br>KO control virus N vs F: $p > 0.9999$<br>KO DREADD virus N vs F: $p = 0.5679$ |
| Figure 2D | two-way ANOVA with Bonferroni | <u>ANOVA</u><br>genotype: $F_{(1,56)} = 25.95$, $p < 0.0001$ |
|  | post hoc comparisons | virus: $F_{(1,56)} = 0.3769$, $p = 0.5417$<br>genotype X virus: $F_{(1,56)} = 0.1461$, $p = 0.7037$<br><u>post hoc comparisons</u><br>WT control virus vs KO control virus: $p = 0.0106$<br>WT DREADD virus vs KO control virus: $p = 0.0012$<br>WT control virus vs KO DREADD virus: $p = 0.0130$<br>WT DREADD virus vs KO DREADD virus: $p = 0.0014$<br>KO control virus vs KO DREADD virus: $p > 0.9999$<br>WT control virus vs WT DREADD virus: $p > 0.9999$ |
| Figure 2E | three-way repeated measures ANOVA with Bonferroni post hoc comparisons | <u>ANOVA</u><br>trial: $F_{(1,56)} = 100.4$, $p < 0.0001$<br>genotype: $F_{(1,56)} = 1.032$, $p = 0.3141$<br>virus: $F_{(1,56)} = 4.179$, $p = 0.0456$<br>trial X genotype: $F_{(1,56)} = 1.731$, $p = 0.1936$<br>trial X virus: $F_{(1,56)} = 14.92$, $p = 0.0003$<br>genotype X virus: $F_{(1,56)} = 3.299$, $p = 0.0747$<br>trial X genotype X virus: $F_{(1,56)} = 5.606$, $p = 0.0214$<br><u>post hoc comparisons</u><br>WT control virus N vs F: $p = 0.0002$<br>WT DREADD virus N vs F: $p < 0.0001$<br>KO control virus N vs F: $p > 0.9999$<br>KO DREADD virus N vs F: $p < 0.0001$ |
| Figure 2F | two-way ANOVA with Bonferroni post hoc comparisons | <u>ANOVA</u><br>genotype: $F_{(1,56)} = 1.731$, $p = 0.1936$<br>virus: $F_{(1,56)} = 14.92$, $p = 0.0003$<br>genotype X virus: $F_{(1,56)} = 5.606$, $p = 0.0214$<br><u>post hoc comparisons</u><br>WT control virus vs KO control virus: $p = 0.0777$<br>WT DREADD virus vs KO control: $p = 0.0039$<br>WT control virus vs KO DREADD virus: $p = 0.4375$<br>WT DREADD virus vs KO DREADD virus: $p > 0.9999$<br>KO control virus vs KO DREADD virus: $p = 0.0002$<br>WT control virus vs WT DREADD virus: $p > 0.9999$ |
| Figure 3C | three-way repeated measures ANOVA with Bonferroni post hoc comparisons | <u>ANOVA</u><br>trial: $F_{(1,58)} = 18.18$, $p < 0.0001$<br>genotype: $F_{(1,58)} = 13.61$, $p = 0.0005$<br>virus: $F_{(1,58)} = 0.03942$, $p = 0.8433$<br>trial X genotype: $F_{(1,58)} = 2.198$, $p = 0.1436$<br>trial X virus: $F_{(1,58)} = 0.09859$, $p = 0.7547$<br>genotype X virus: $F_{(1,58)} = 0.1478$, $p = 0.7021$<br>trial X genotype X virus: $F_{(1,58)} = 1.528$, $p = 0.2214$<br><u>post hoc comparisons</u><br>WT control virus N vs F: $p = 0.0270$<br>WT DREADD virus N vs F: $p = 0.3566$ |
| | | KO control virus N vs F: $p > 0.9999$<br>KO DREADD virus N vs F: $p = 0.3094$ |
| Figure 3D | two-way ANOVA<br>with Bonferroni<br>post hoc<br>comparisons | <u>ANOVA</u><br>genotype: $F_{(1,58)} = 2.198$, $p = 0.1436$<br>virus: $F_{(1,58)} = 0.09859$, $p = 0.7547$<br>genotype X virus: $F_{(1,58)} = 1.528$, $p = 0.2214$<br><u>post hoc comparisons</u><br>WT control virus vs KO control virus: $p = 0.3322$<br>WT DREADD virus vs KO control: $p > 0.9999$<br>WT control virus vs KO DREADD virus: $p > 0.9999$<br>WT DREADD virus vs KO DREADD virus: $p > 0.9999$<br>KO control virus vs KO DREADD virus: $p > 0.9999$<br>WT control virus vs WT DREADD virus: $p > 0.9999$ |
| Figure 3E | three-way<br>repeated<br>measures<br>ANOVA with<br>Bonferroni post<br>hoc comparisons | <u>ANOVA</u><br>trial: $F_{(1,58)} = 40.21$, $p < 0.0001$<br>genotype: $F_{(1,58)} = 0.04443$, $p = 0.8338$<br>virus: $F_{(1,58)} = 1.340$, $p = 0.2518$<br>trial X genotype: $F_{(1,58)} = 3.169$, $p = 0.0803$<br>trial X virus: $F_{(1,58)} = 3.339$, $p = 0.0728$<br>genotype X virus: $F_{(1,58)} = 1.233$, $p = 0.2715$<br>trial X genotype X virus: $F_{(1,58)} = 4.317$, $p = 0.0422$<br><u>post hoc comparisons</u><br>WT control virus N vs F: $p = 0.0021$<br>WT DREADD virus N vs F: $p = 0.0071$<br>KO control virus N vs F: $p > 0.9999$<br>KO DREADD virus N vs F: $p = 0.0004$ |
| Figure 3F | two-way ANOVA<br>with Bonferroni<br>post hoc<br>comparisons | <u>ANOVA</u><br>genotype: $F_{(1,58)} = 3.169$, $p = 0.0803$<br>virus: $F_{(1,58)} = 3.339$, $p = 0.0728$<br>genotype X virus: $F_{(1,58)} = 4.317$, $p = 0.0422$<br><u>post hoc comparisons</u><br>WT control virus vs KO control virus: $p = 0.0446$<br>WT DREADD virus vs KO control: $p = 0.0827$<br>WT control virus vs KO DREADD virus: $p > 0.9999$<br>WT DREADD virus vs KO DREADD virus: $p > 0.9999$<br>KO control virus vs KO DREADD virus: $p = 0.0268$<br>WT control virus vs WT DREADD virus: $p > 0.9999$ |
| Figure 4C | three-way<br>repeated<br>measures<br>ANOVA with<br>Bonferroni post<br>hoc comparisons | <u>ANOVA</u><br>drug: $F_{(1,113)} = 4.868$, $p = 0.0294$<br>genotype: $F_{(1,113)} = 5.530$, $p = 0.0204$<br>virus: $F_{(1,113)} = 0.2183$, $p = 0.6412$<br>drug X genotype: $F_{(1,113)} = 2.812$, $p = 0.0963$<br>drug X virus: $F_{(1,113)} = 0.1906$, $p = 0.6633$<br>genotype X virus: $F_{(1,113)} = 0.9545$, $p = 0.3307$<br>drug X genotype X virus: $F_{(1,113)} = 4.944$, $p = 0.0282$ |
|  |  | <u>post hoc comparisons</u><br>WT control virus vehicle vs CNO: $p > 0.9999$<br>WT DREADD virus vehicle vs CNO: $p > 0.9999$<br>KO control virus vehicle vs CNO: $p > 0.9999$<br>KO DREADD virus vehicle vs CNO: $p = 0.0147$ |
| Figure 4E | three-way repeated measures ANOVA with Bonferroni post hoc comparisons | <u>ANOVA</u><br>drug: $F_{(1,113)} = 3.685, p = 0.0574$<br>genotype: $F_{(1,113)} = 0.1695, p = 0.6813$<br>virus: $F_{(1,113)} = 0.7422, p = 0.3908$<br>drug X genotype: $F_{(1,113)} = 0.1344, p = 0.7146$<br>drug X virus: $F_{(1,113)} = 0.9878, p = 0.3224$<br>genotype X virus: $F_{(1,113)} = 4.740, p = 0.0316$<br>drug X genotype X virus: $F_{(1,113)} = 0.3930, p = 0.5320$<br><u>post hoc comparisons</u><br>WT control virus vehicle vs CNO: $p > 0.9999$<br>WT DREADD virus vehicle vs CNO: $p > 0.9999$<br>KO control virus vehicle vs CNO: $p = 0.7543$<br>KO DREADD virus vehicle vs CNO: $p > 0.9999$ |
| Figure 4G | three-way repeated measures ANOVA with Bonferroni post hoc comparisons | <u>ANOVA</u><br>drug: $F_{(1,113)} = 3.624, p = 0.0595$<br>genotype: $F_{(1,113)} = 0.2965, p = 0.5872$<br>virus: $F_{(1,113)} = 0.7946, p = 0.3746$<br>drug X genotype: $F_{(1,113)} = 3.191, p = 0.0767$<br>drug X virus: $F_{(1,113)} = 0.6312, p = 0.4286$<br>genotype X virus: $F_{(1,113)} = 0.3050, p = 0.5819$<br>drug X genotype X virus: $F_{(1,113)} = 0.1184, p = 0.7314$<br><u>post hoc comparisons</u><br>WT control virus vehicle vs CNO: $p > 0.9999$<br>WT DREADD virus vehicle vs CNO: $p > 0.9999$<br>KO control virus vehicle vs CNO: $p = 0.2643$<br>KO DREADD virus vehicle vs CNO: $p > 0.9999$ |

**Supplementary Table 2.** Detailed statistics for supplementary figures.

| Data set | Type of test | Confidence |
| --- | --- | --- |
| Supplementary Figure 1B | unpaired t-test | Open: $t_{38} = 0.992$, $p = 0.327$<br>Closed: $t_{38} = 2.016$, $p = 0.0509$<br>Center: $t_{38} = 7.485$, $p < 0.0001$ |
| Supplementary Figure 1C | unpaired t-test | Open: $t_{38} = 0.841$, $p = 0.406$<br>Closed: $t_{38} = 3.548$, $p = 0.0011$ |
| Supplementary Figure 2A | unpaired t-test | $t_{18} = 2.265$, $p = 0.0360$ |
| Supplementary Figure 2B | unpaired t-test | $t_{18} = 0.831$, $p = 0.416$ |
| Supplementary Figure 3A | Mann-Whitney U test | $U_{16} = 21.50$ $p = 0.0984$ |
| Supplementary Figure 3B | unpaired t-test | $t_{16} = 0.5054$, $p = 0.6202$ |
| Supplementary Figure 4A | three-way repeated measures ANOVA with Bonferroni post hoc comparisons | <u>ANOVA</u><br>drug: $F_{(1,113)} = 2.880$, $p = 0.0924$<br>genotype: $F_{(1,113)} = 3.389$, $p = 0.0682$<br>virus: $F_{(1,113)} = 1.076$, $p = 0.3019$<br>drug X genotype: $F_{(1,113)} = 1.851$, $p = 0.1764$<br>drug X virus: $F_{(1,113)} = 0.5778$, $p = 0.4487$<br>genotype X virus: $F_{(1,113)} = 3.940$, $p = 0.0496$<br>drug X genotype X virus: $F_{(1,113)} = 3.264$, $p = 0.0735$<br><u>post hoc comparisons</u><br>WT control virus vehicle vs CNO: $p > 0.9999$<br>WT DREADD virus vehicle vs CNO: $p > 0.9999$<br>KO control virus vehicle vs CNO: $p = 0.0947$<br>KO DREADD virus vehicle vs CNO: $p > 0.9999$ |
| Supplementary Figure 4B | three-way repeated measures ANOVA with Bonferroni post hoc comparisons | <u>ANOVA</u><br>drug: $F_{(1,113)} = 0.007624$, $p = 0.9305$<br>genotype: $F_{(1,113)} = 0.0007800$, $p = 0.9778$<br>virus: $F_{(1,113)} = 4.221$, $p = 0.0422$<br>drug X genotype: $F_{(1,113)} = 2.757$, $p = 0.0996$<br>drug X virus: $F_{(1,113)} = 9.524$, $p = 0.0026$<br>genotype X virus: $F_{(1,113)} = 0.3959$, $p = 0.5305$<br>drug X genotype X virus: $F_{(1,113)} = 0.04674$, $p = 0.8292$<br><u>post hoc comparisons</u><br>WT control virus vehicle vs CNO: $p > 0.9999$<br>WT DREADD virus vehicle vs CNO: $p = 0.3150$<br>KO control virus vehicle vs CNO: $p = 0.2008$<br>KO DREADD virus vehicle vs CNO: $p > 0.9999$ |

## Notes

### Competing Interest Statement

The authors have declared no competing interest.

